# Different evolutionary pathways of HIV-1 between fetus and mother perinatal transmission pairs indicate unique immune selection pressure in fetuses

**DOI:** 10.1101/2020.08.28.272930

**Authors:** Manukumar Honnayakanahalli Marichannegowda, Michael Mengual, Amit Kumar, Elena E. Giorgi, Joshua J. Tu, David R. Martinez, Xiaojun Li, Liping Feng, Sallie R Permar, Feng Gao

**Affiliations:** Department of Medicine, Duke University Medical Center, Durham, NC 27710, USA; Duke Human Vaccine Institute, Duke University Medical Center, Durham, NC 27710, USA; Theoretical Division, Los Alamos National Laboratory, Los Alamos, NM 87544, USA; Department of Molecular Genetics and Microbiology, Duke University Medical Center, Durham, NC 27710, USA; Department of Pediatrics, Duke University Medical Center, Durham, NC 27710, USA; Department of Obstetrics and Gynecology, Duke University Medical Center, Durham, NC 27710, USA

**Author notes:** These authors contributed equally (Manukumar Honnayakanahalli Marichannegowda, Michael Mengual). These authors equally supervised this work (Sallie R Permar, Feng Gao).

**Keywords:** HIV-1, Mother-to-infant-transmission, Envelope glycoprotein, Fetus, *In utero*, Selection

## Abstract

Study of evolution and selection pressure on HIV-1 in fetuses will lead to a better understanding of the role of immune responses in shaping virus evolution and vertical transmission. Detailed genetic analyses of HIV-1 *env* gene from 12 *in utero* transmission pairs show that most infections (67%) occur within two months from childbirth. In addition, e*nv* sequences from long-term infected fetuses were highly divergent and formed separate phylogenetic lineages from their cognate maternal viruses. Host selection sites unique to infant viruses were identified in regions frequently targeted by broadly neutralizing antibodies and T cell immune responses. The identification of unique selection sites in the *env* gene of fetal viruses indicates that the immune system in fetuses is capable of exerting selection pressure on viral evolution. Studying selection and evolution of HIV-1 or other viruses in fetuses can be an alternative approach to investigate adaptive immunity in fetuses.

## Introduction

Among 38 million people currently living with HIV/AIDS, 1.8 million are children infected through mother-to-child transmission (MTCT) (UNAIDS, 2020). Perinatal HIV-1 infection can be acquired through antepartum (*in utero*), intrapartum (around delivery), or postpartum (through breast-feeding). Prior to the availability of antiretroviral therapy (ART), the rate of HIV-1 MTCT among pregnant women was about 30-40% (Wolinsky et al., 1992) and it is estimated that each transmission route accounted for about one-third of the total MTCT transmissions. After the introduction of ART, the MTCT rate was significantly reduced to ~2% (Dorenbaum et al., 2002; Mahy et al., 2010). However, MTCT still leads to well over 100,000 new infant infections annually. With the “90-90-90” campaign by the United Nations Program on HIV/AIDS (UNAIDS), it was expected that pediatric HIV-1 infections would be reduced to fewer than 20,000 by 2020 (Bailey et al., 2018). However, since HIV testing and ART are not widely accessible to all pregnant women in resource-limited countries, this goal will not be achieved (Kellerman and Essajee, 2010; Tang and Nour, 2010). Further, as ART is often initiated upon diagnosis of HIV-1 during prenatal visits or around the time of delivery, *in utero* infections are responsible for a higher portion of the ongoing cases of MTCT than would have been the case in the pre-ART era.

During pregnancy, passive transfer of maternal IgG antibodies (Abs) to fetuses may play a role in blocking MTCT and exert immune selection pressure on viruses in fetuses (Albrecht and Arck, 2020; Fouda et al., 2018; Martinez et al., 2017; Permar et al., 2015), but the mechanism is still poorly understood (Albrecht and Arck, 2020; Martinez et al., 2019; Palmeira et al., 2012; Wilcox et al., 2017). The relatively high rate of *in utero* MTCT suggests that transferred maternal Abs cannot fully prevent HIV-1 infection of fetuses, possibly due to the presence of escape variants resistant to maternal neutralizing antibodies (nAbs). Our previous studies showed that the transmitted viruses in infants are often resistant to neutralization by cognate maternal plasma (Kumar et al., 2018; Martinez et al., 2020; Martinez et al., 2017; Moody et al., 2015; Permar et al., 2015), yet it remains unclear if maternal antibodies contribute to blocking placental HIV transmission.

Innate immune responses and antigen-specific T cell responses to pathogens are detectable in fetuses, suggesting exposure to maternal infection *in utero* may prime the developing immune system (Wilcox and Jones, 2018). In fact, aberrant fetal immune responses to maternal antigens may play an important role in the pathogenesis of preterm labor (Frascoli et al., 2018). A recent study demonstrated that fetal antigen presenting cells (APC), such as conventional dendritic cells (DC), can respond to tolllike receptors (TLR) as early as the second trimester (McGovern et al., 2017). However, a comparison of immune responses between fetuses and infants shows that the immune responses mediated by macrophages are more geared towards immune tolerance in fetuses. Little is known about the function of fetal B lymphocytes in response to infectious agents (Dauby and Marchant, 2020). Fetal exposure to HIV-1 primes the immune system to enhance immune activation and alter T cell homing in fetuses (Bunders et al., 2014). These studies indicate that innate and adaptive immune responses are functioning in the fetus, but none have demonstrated that immune responses can exert selection pressure on viral evolution during *in utero* infection.

HIV-1 infection of fetuses can occur in pregnant women without ART, but the timing and frequency of infection as well as viral evolution in fetuses remains unknown. Moreover, the role of immune selection on the viruses in fetuses has not yet been established. To address these questions, we have studied a total of 721 HIV-1 *env* gene sequences from 12 pairs of mothers and infants who were infected before delivery. We found that most *in utero* infections occur late in pregnancy and unique strong selection on viral evolution during long-term *in utero* infection is different between fetuses and their cognate mothers, suggesting the presence of adaptive immune responses against HIV-1 in fetuses.

## Materials and methods

### Sample collection

Plasma samples were collected from 10 pairs of mothers and infants from the Women Infant Transmission Study (WITS) cohort (Martinez et al., 2017; Permar et al., 2015) and 2 pairs from the CHAVI009 cohort (Moody et al., 2016). All infants were diagnosed as *in utero* infection by positive PCR at birth using genomic DNA extracted from peripheral blood mononuclear cells from the newborn babies. Plasma samples were collected from infants between delivery and 54 days after birth and from mothers at delivery. Samples from two different time points were collected from four infants (1039i, 2564i, 1348i and 1580i). Two mothers (9112m and 3915m) from the CHAVI009 cohort received 1 dose of nevirapine at delivery. No other individual was on antiretroviral therapy.

Placental plasma was obtained as described previously from the CHAVI009 cohort (Anderson, 1997). Briefly, an incision was made at the cleaned maternal surface of the placenta, and blood from the incision was aspirated into tubes containing EDTA (Ethylenediaminetetraacetic acid), sodium citrate and heparin. The samples were then separated and the plasma was stored at −80°C.

### Ethics statement

All samples in the study used were received from the existing cohorts WITS and CHAVI09, which are deidentified and deemed as research not involving human subjects by Duke University Institutional Review Board. The protocol number is Pro00016627.

### Analysis of the *env* gene sequences by single genome amplification

Viral RNA was extracted from plasma samples using PureLink Viral RNA/DNA mini kit (Invitrogen, Carlsbad, CA) and used to generate complementary DNA (cDNA) using SuperScript III reverse transcription mix (Invitrogen) and antisense primer R3.B3R (5’-ACTACTTGAAGCACTCAAGGCAAGCTTTATTG-3’; according to nucleotide [nt] position 9611-9642 in HXB2) for the 3’-half genome. The synthesized cDNA was subjected to single genome amplification (SGA) analysis using Platinum Taq DNA polymerase High Fidelity (Invitrogen) (Kumar et al., 2018). The first round PCR was performed using the primers 07For7 (5’-AAATTAYAAAAATTCAAAATTTTCGGGTTTATTACAG-3’; nt 4875-4912) and 2.R3.B6R (5’-TGAAGCACTCAAGGCAAGCTTTATTGAGGC-3’; nt 9636–9607), and second round PCR was carried out using 2μl of the first round product with primers VIF1 5’-GGGTTTATTACAGGGACAGCAGAG-3’; nt 5960–5983) and Low2c 5’-TGAGGCTTAAGCAGTGGGTTCC-3’; nt 9413–9436). The first round PCR conditions were as follows: one cycle at 94 °C for 2 min; 35 cycles of a denaturing step at 94 °C for 15 s, an annealing step at 58 °C for 30 s, an extension step at 68 °C for 4 min; and one cycle of an additional extension at 68 °C for 10 min. The second round PCR conditions were the same as for the first round PCR, except that 45 thermocycling cycles were used. The final PCR products were visualized by agarose gel electrophoresis, purified and sequenced for the entire *env* gene using ABI3730xl genetic analyzer (Applied Biosystems, Foster City, CA).

### Sequence analysis

Each SGA amplicon was sequenced using the primer walking method and all sequences were assembled and edited using Sequencher 4.5 (Gene Codes, Ann Arbor MI). The final assembled sequences from each infant and mother transmission pair were aligned using the Gene Cutter tool (http://www.hiv.lanl.gov/content/sequence/GENE_CUTTER/cutter.html), and the alignments were optimized manually using Seaview (Gouy et al., 2010). The Highlighter plots were generated using the Highlighter tool (https://www.hiv.lanl.gov/content/sequence/HIGHLIGHT/highlighter_top.html).

APOBEC-mediated enrichment of G-to-A substitutions was screened using the Hypermut tool (http://www.hiv.lanl.gov/content/sequence/HYPERMUT/hypermut.html) and sequences with significant hypermutation (p<0.1) were excluded from further analysis. Recombinants were detected using the RAPR tool (https://www.hiv.lanl.gov/content/sequence/RAP2017/rap.html). Recombinants and their putative descendants were excluded from the timing analysis. Neighbor-joining trees were constructed using the Kimura 2-parameter model with 1000 bootstrap replications. Phylogenetic trees and highlighter plots were used to infer the infant T/F *env* gene sequences as previously described (Keele et al., 2008). Infection time was estimated using the Poisson Fitter tool (http://www.hiv.lanl.gov/content/sequence/POISSONFITTER/pfitter.html) after excluding all hypermutated and recombinant sequences (Giorgi et al., 2010). Time estimates for infants with multiple infecting strains were obtained on each individual lineage and then averaged using a harmonic mean. For two long-term infected infants, 1039i and 1580i, we obtained a lower boundary on the number of days since the most recent common ancestor (MRCA) by separating into sublineages for the remaining viral population after removing all detected recombinants. In infant 1039i, the remaining population formed a single lineage that showed evidence of non-random accumulation of mutations at three sites in particular, 404, 421, and 566. Therefore, in order to infer the time since the selection bottleneck, those sites were masked for the Poisson Fitter analysis. Infant 1580i, on the other hand, did not show any evidence of selection sites after removing all recombinants. The remaining viral population could be divided into three sublineages, of which two had originated through recombination events identified by the RAPR analysis.

Within-lineage genetic pairwise *p*-distances were calculated using sequence data from each sample, after removing recombinants and hypermutated sequences, and compared between infant and maternal viral populations. The accumulation of synonymous and non-synonymous mutations across the entire *env* gene was determined by pairwise comparison of all the *env* sequences in infants or mothers according to the method developed by Nei and Gojobori (Nei and Gojobori, 1986) using SNAP (https://www.hiv.lanl.gov/content/sequence/SNAP/SNAP.html). Sites under significant diversifying selection pressure were identified using the MEME tool, available on the datamonkey server (Murrell et al., 2012). The GenBank accession numbers for the sequences from this study are MT853116 to MT853181 and MT861212-MT861991.

### Generation of Env-pseudoviruses

Two *env* genes that represent the consensus sequences (infant 1 and 2) of two sub-clusters in the viral population in infant 1580i and one *env* gene (M.2) that represents one major cluster viruses in mother 1580m were codon optimized and chemically synthesized (Thermo Fisher, Waltham, MA, USA). The *env* genes were cloned into pcDNA3.1(+) at the Hind III and EcoR I sites. The selection of MTCT pair 1580 is because it is the only pair from which the plasma samples from both the infant and the mother were available for autologous neutralization analysis. The *env* gene clones together with the *env*-deficient HIV-1 plasmid DNA (SG3Δenv) were contransfected into HEK293T cells (ATCC, Manassas, VA, USA) using the FuGene 6 transfection reagent (Promega, Madison, WI, USA). Two days after transfection, the culture supernatant containing pseudoviruses was harvested, aliquoted, and stored at −80°C.

### Neutralization Assay

Neutralization activities of the plasma samples were determined by the single-round infection of HIV-1 Env-pseudoviruses in TZM-bl cells as described previously (Montefiori, 2009). Briefly, plasma samples were first heat-inactivated by incubating at 56°C for 60 min. After the 1:3 serial diluted plasma samples were incubated with Env-pseudoviruses at 37°C for 1 hour, the mixtures were used to infect TZM-bl cells. Plates were read after incubating at 37 °C for 48 hours. The 50% inhibitory dose (ID_50_) was defined as plasma reciprocal dilution at which relative luminescence units (RLU) were reduced by 50% compared with RLU in virus control wells after subtraction of background RLU in cell control wells. A response was considered positive if the neutralization titer was higher than 1:30.

### Detection of HIV-1 specific antibodies by ELISA

HIV-1 specific binding antibody titers were performed by coating high-binding 384 well plates (Corning, Corning, NY) overnight at 4°C with 1086C and Con6 gp120s. Plates were washed once and blocked for 1 hour at room temperature (RT) with SuperBlock (4% whey protein, 15% goat serum, and 0.5% Tween 20 diluted in 1X phosphate-buffered saline [PBS]). Plates were washed and 10 μl of plasma in 3-fold serial dilutions were added in duplicate and incubated at room temperature for 1 hour. Plates were washed two times and horseradish peroxidase (HRP)-conjugated goat anti-human IgG antibody (Sigma Aldrich, St. Louis, MO) was used at a 1:5,000 dilution and incubated at room temperature for 1 hour. Plates were washed four times and SureBlue reserve TMB substrate (KPL, Gaithersburg, MD) was added. Reactions were stopped by stop solution (KPL, Gaithersburg, MD) and optical densities (OD) were measured at 450nm. Concentrations were calculated using a VRC01 IgG standard curve (at a 2-fold series from 0.0005 to 1 μg/mL).

### Depletion of IgG in plasma

IgG depletion was performed by diluting systemic blood and placenta plasma by 2-fold with Tris-buffered saline (TBS) and filtering in Spin-X centrifuge tube filters (Corning). Diluted and filtered samples were then added to a Protein G High Performance MultiTrap 96-well plate (GE Healthcare, Chicago, IL) and shaken at 1,100 RPM for one hour at room temperature. IgG-depleted plasma samples were obtained by centrifugation at 700g for 3 minutes onto a receiving plate.

### Quantification and statistical analysis

Sites under significant diversifying selection pressure were identified using the MEME tool, available on the datamonkey server (Murrell et al., 2012). All statistical comparisons of p-distances and the rate of nonsynonymous substitutions per nonsynonymous site (dN/dS) were conducted in R (R Development Core Team, 2009).

## Results

### Most *in utero* MTCTs occur during the late stage of pregnancy

Twelve *in utero* mother-infant transmission pairs were selected from two cohorts (10 from WITS and 2 from CHAVI09). All infants were infected *in utero* as determined by positive PCR with HIV-1 DNA genome extracted from infant blood samples at birth. All but one mother (0842) had plasma samples available at delivery. Infant plasma samples were available from various time points after delivery (3 at delivery, 7 at ~1 month after birth, 2 at ~2 months after birth). A total of 721 HIV-1 *env* gene sequences (317 and 404 from mothers and infants, respectively) were obtained by single genome amplification (SGA) (**Table 1**). All sequences from each mother-infant pair were clustered together in the overall phylogenetic tree, and sequences from different mother-infant pairs were phylogenetically distinctive from each other. Phylogenetic tree and Highlighter plot analyses showed that the *env* sequences from five infants (0782i, 1730i, 0489i, 1348i and 9112i) were highly homogeneous and a single transmitted/founder (T/F) viral sequence was inferred for each infant (**Figs. 1 and S1**). The *env* sequences from three infants were heterogenous, consisting of two or three homogenous sub-populations. Among them, two T/Fs were inferred for two infants (1744i and 0842i) and three T/Fs for the third infant (3915i). The *env* sequences from four other infants (1039i, 2093i, 1580i, and 2564i) were highly heterogenous and no T/F sequences could be reliably inferred (**Figs. 1 and S1**). Of note, these four infants were on average sampled at an older age: the mean number of days after birth was 43 days, compared to 17 days for the other 8 infants.

**Table 1.** Demographic and genetic characteristics of mother-infant transmission pairs

| transmission pair | Viral load (copies/ml) |  | No. of SGAs |  | No. of T/Fs | Infant age (days) | Estimated transmission time in days (95%, CI) | Transmission days from birth |
| --- | --- | --- | --- | --- | --- | --- | --- | --- |
|  | Mother | Infant | Mother | Infant |  |  |  |  |
| 0782 | 8,224 | 2,716 | 32 | 7 | 1 | 6 | 9 (4,15) | -3 |
| 1730 | 45,924 | 716,003 | 28 | 29 | 1 | 27 | 44 (33,54) | -17 |
| 0489 | 30,067 | 779,602 | 35 | 26 | 1 | 20 | 32 (21,44) | -12 |
| 1348 | 4,458 | 332,727 | 19 | 46 | 1 | 40 | 61 (52,70) | -21 |
| 9112 | NA | NA | 34 | 21 | 1 | 1 | 20 (14,26) | -19 |
| 1744 | 276,955 | NA | 21 | 35 | 2 | 21 | 49 (29,69) | -28 |
| 0842 | 6,057 | 216,000 | NA | 37 | 2 | 20 | 27 (23,32) | -7 |
| 3915 | NA | NA | 41 | 51 | 3 | 1 | 38 (35,41) | -37 |
| 1039 | 25,049 | 1,957,418 | 24 | 30 | NR | 39 | ND |  |
| 2093 | 3,101,258 | 1,119,972 | 38 | 45 | NR | 54 | ND |  |
| 1580 | 78,174 | NA | 32 | 53 | NR | 51 | ND |  |
| 2564 | 26,640 | NA | 13 | 24 | NR | 28 | ND |  |

Transmission time was estimated using the LANL Poisson Fitter tool.
NA: Not available.
NR: Number of T/Fs was not retrievable for 4 chronically infected infants due to the high genetic diversity of the viral population.
ND: Time of infection is not determined.

**Fig. 1.**
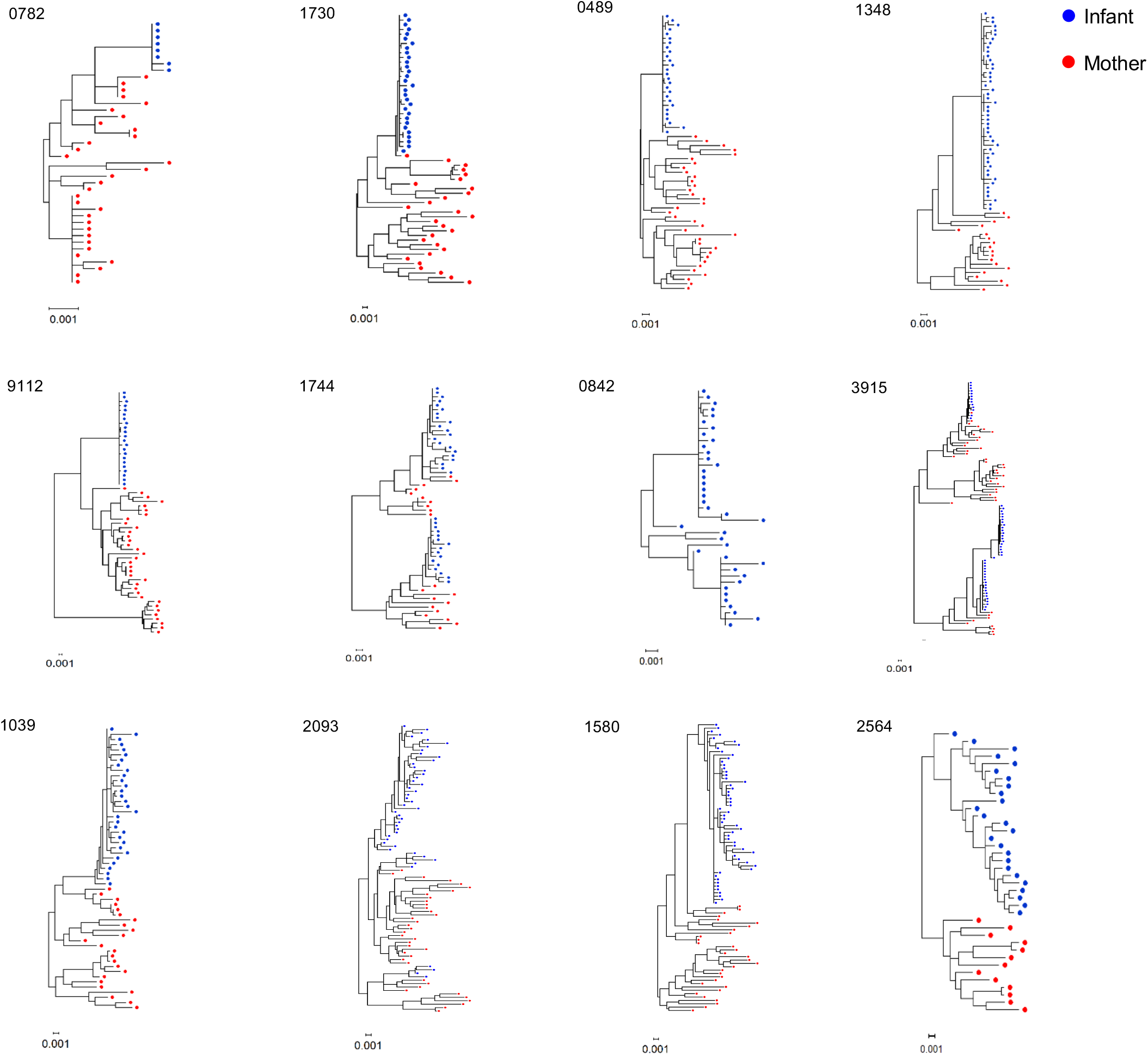
Phylogenetic trees of the *env* sequences from each mother-infant transmission pair. Phylogenetic trees were constructed using the neighbor-joining method with the Kimura 2-parameter model. Sequences from infants and mothers are indicated in blue and red dots, respectively.

Using the Poisson Fitter tool (Giorgi et al., 2010), we were able to determine the infection time for the eight infant samples with well-defined T/F lineages, which ranged from 9 to 61 days, corresponding to 3-21 days before birth (**Table 1**). For infants 1744i, 0842i and 3915i, who were infected with 2 or 3 distinct T/F viruses, the overall time since infection was calculated by taking the harmonic mean of the time estimates from each separate lineage, as previous described (Giorgi et al., 2010). Their infection times were 49, 27 and 38 days, respectively, corresponding to 28, 7 and 37 days before birth, respectively. The estimated *in utero* infection time for 0782i and 0842i were the shortest, only 3 and 7 days before birth, respectively. Importantly, all estimated times of infection were before their corresponding birth dates, further confirming that all infants had been infected *in utero*, within 40 days before birth (**Table 1**).

The *env* sequences in four infants (1039i, 2093i, 1580i and 2564i) were distinctive from all other infants in that their genetic diversity was far more divergent (**Figs. 1 and S1**), indicating that these infections were too far out to allow the use the Poisson Fitter tool, which is based on the assumption of random accumulation of mutations prior to the onset of selection. The BEAST (Bayesian evolutionary analysis by sampling trees) software package has been widely used to estimate the time of the most recent common ancestor (MRCA) or infection time (Suchard et al., 2018). However, in this context, BEAST is also limited for two reasons. First, any timing inference on the infant sequences yields the time since the MRCA, which likely took place in the mother, not the infant. Indeed, when we ran BEAST on these four infants, we obtained timing estimates far longer than the 40-week time period of pregnancy. Second, further analyses using the recombination detection tool RAPR (Song et al., 2018) reveal extensive recombination in all four infant samples (**Fig. S2A**). This degree of pervasive recombination can cause branch length artifacts in the phylogenetic reconstructions upon which the BEAST methods are based, potentially confounding the time estimates (Giorgi et al., 2020).

Indeed, the RAPR analysis revealed that for two infants in particular, 2564i and 2093i, all but 3 or less sequences had derived from recombination events. In our previous study, we showed that in heterogeneous infections (multiple T/Fs), recombinants rapidly replace the lineages evolving from the initial infecting strains, with a median half-time of 27 days (Song et al., 2018). This result is consistent with the two infants being infected *in utero* with multiple TFs.

For the other two infants (1039i and 1580i), after removing all recombinants and their descendants, there were enough sequences left to discern lineages (**Fig. S2B**) and infer infection times using our previous methods. While we cannot discern with certainty whether these lineages are the original T/Fs or whether, more likely given the level of accumulated diversity in the original sample, are in fact subclades arose from subsequent bottlenecks, the timing of these sequences provides useful information on the least number of days since the MRCA. The minimum number of days since infection were 104 (95% CI 84-123 days) for infant 1039i after removing recombinants and selection epitopes, and 53 (95% CI 29-76 days) for infant 1580i based on the “oldest” lineage (see Materials and Methods).

Taken together, these results show that the majority of *in utero* infections (67%) occur during the last two months of the pregnancy in this study.

### High genetic diversity of viral populations during long-term *in utero* infection

To determine the genetic diversity accumulated during *in utero* infections, we calculated the genetic p-distances for the viral population in each infant and compared the diversity levels of viruses in the infants to their cognate mothers. The within-lineage viral populations in eight infants from whom the T/F viral sequences could be inferred were highly homogenous. As expected, the average diversity of the within-lineage viral populations from each T/F virus was much lower in the infants (0.12%; 0.02%-0.27%) than in the mothers (1.56%; 0.27%-3.23%) (**Fig. 2**). However, the genetic diversity levels were much higher in viral populations of the four infants from whom the T/F viral sequences could be not be inferred, which is also to be expected given the high level of recombination detected in these samples. In these infants, the viral population average diversity was 0.62% (0.27%-1.13%), compared to the 3-fold higher diversity observed in their cognate maternal viral populations (1.68%; 1.3%-2.39%). While the diversity levels were similar among the maternal viruses from both groups (1.56% vs. 1.68%), the diversity of the viral population in long-term infection infants was 5 times higher than that in short-term infection infants. In pair 2564, the average diversity (0.75%) in 2564i was less than two-fold difference of that (1.3%) in 2564m (**Fig. 2**). However, the p-diversity values in the short-term infections were calculated within lineages after removing all detected recombinants. Lineages in two of the long-term infected infants were no longer detectable, but, for the two infants for which tentative lineages were identified after removing all possible recombinant sequences, the average p-diversity was 0.38% and 0.35%, which were a little less than half of diversities when recombinant sequences were included for analysis, but still 3-folds higher than the mean diversity in the infants with clearly defined T/F sequences. This indicates that the recombination played an important role in increased diversity in these long-term infected infants.

**Fig. 2.**
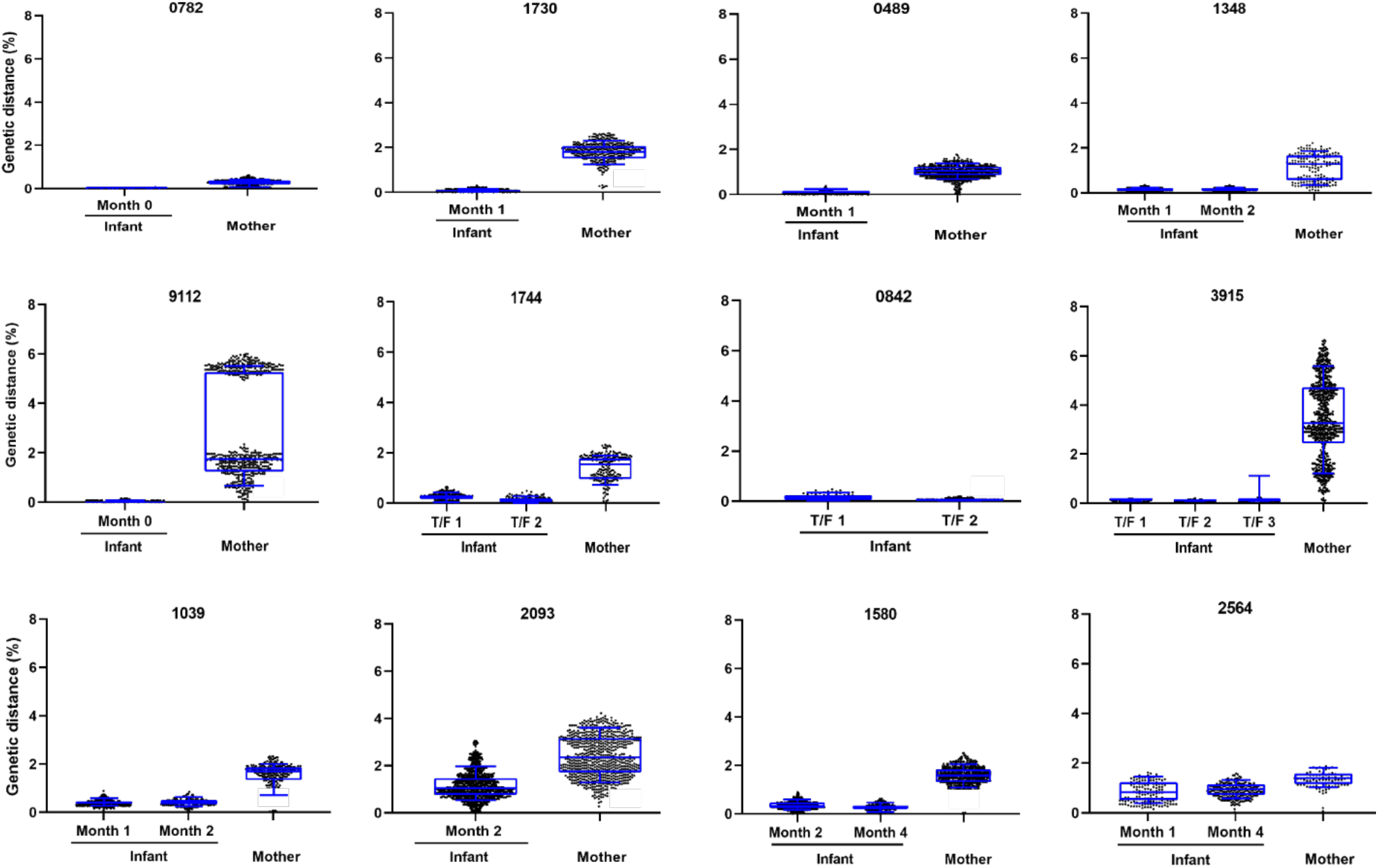
Comparison of genetic diversity of the viral populations of mother and infant in each transmission pair. Genetic diversity was measured by calculating within lineage pairwise genetic distances (p-distance) of maternal sequences and infant sequences at different time points. P-distances are plotted as black dots, and their median in blue lines. The middle blue line indicates the median diversity, box shows 25 to 75 percentiles of the data, and whisker shows 10 to 90 percentiles of the data.

One caveat with the higher diversity observed in the four long-term infected infants is that these infants were on average sampled at a later time (on average, 43 days after birth) than the other eight infants (on average, 17 days after birth). The one exception is infant 1348i, sampled at 40 days after birth, whose infection was clearly initiated by a single T/F virus. While recombination is pervasive in all HIV infections, due to the homogeneity of the sequences, it takes longer for recombinants to be detectable in a single T/F infection (Song et al., 2018). Therefore, it is likely that the long in utero infection time allows transmission of multiple T/F viruses to the fetuses and extensive viral recombination to take place, so that, by the time these infants were sampled, recombinants had completely replaced the infecting viral populations.

Additionally, infant 1348i, together with three long-term infected infants 1039i, 1580i, and 2564i, were all sampled a second time (**Figure S3A**), one to three months later (35 days later for 1348i; 26, 51 and 85 days later for 1039i, 1580i, and 2564i, respectively), and all showed no significant increase in p-diversity from the first time point (p=0.9 by paired Wilcoxon test; Fig. 2) and they were phyloggeneitcaly indistinguishable (**Figure S3B**). These results indicate that the viral genetic diversity generally remained similar in infants after additional 1-3 months of infection after birth, demonstrating that the mutations accumulated after birth only marginally contributed to the high levels of genetic variation of viruses in those long-term *in utero* infections. This suggests that these viruses are under host selection pressure while replicating *in utero*.

### Transmission of multiple T/F viruses in long-term *in utero* infections

The amino acid sequences in the infant viral populations present many unique divergent sequence motifs in the *env* genes. We sought to investigate whether these sequence motifs observed in the infants had originated in their cognate mothers or evolved in fetuses. Two predominant sequence motifs (A and B) and four unique sequence motifs in variable loop 1 (V1) were detected in 2093i (**Fig. 3**). Sequences with identical motif A and nearly identical motif B in the infant were also found in the cognate maternal viruses (represented by 1 and 3 sequences, respectively). Motif C, represented by a single sequence (2093i.5), was found in two of the mother’s viral sequences. In the V3/C3/V4 region, there were two sequence motifs in the infant viral population (D and E, **Fig. 3**). The minority motif D was found in three infant sequences and, a similar form was found in one maternal sequence (2093m.20). Additionally, two more unique motifs were found in the infant (sequences 2093i. 18 and 2093i.5a, highlighted in red and gray in Fig. 3). Interestingly, a motif identical to the one found in 2093i.18 (highlighted in red) was also found in one maternal viral sequence (2093m.26) and motifs similar to the ones in 2093i.5a (highlighted in grey) were detected in four maternal viral sequences (2093m.5, 2093m.31, 2093m.38s and 2093m.42). No sequences in the mother were found to be identical to the predominant motif E sequence in the infant, although one maternal viral sequence (2093m.3) was very similar. Infant sequences carrying motifs similar to the ones observed in the maternal viral population indicate that these viruses most likely derived from the transmitted viruses from the cognate mother viral population, rather than emerging *de novo* in the infant.

**Fig. 3.**
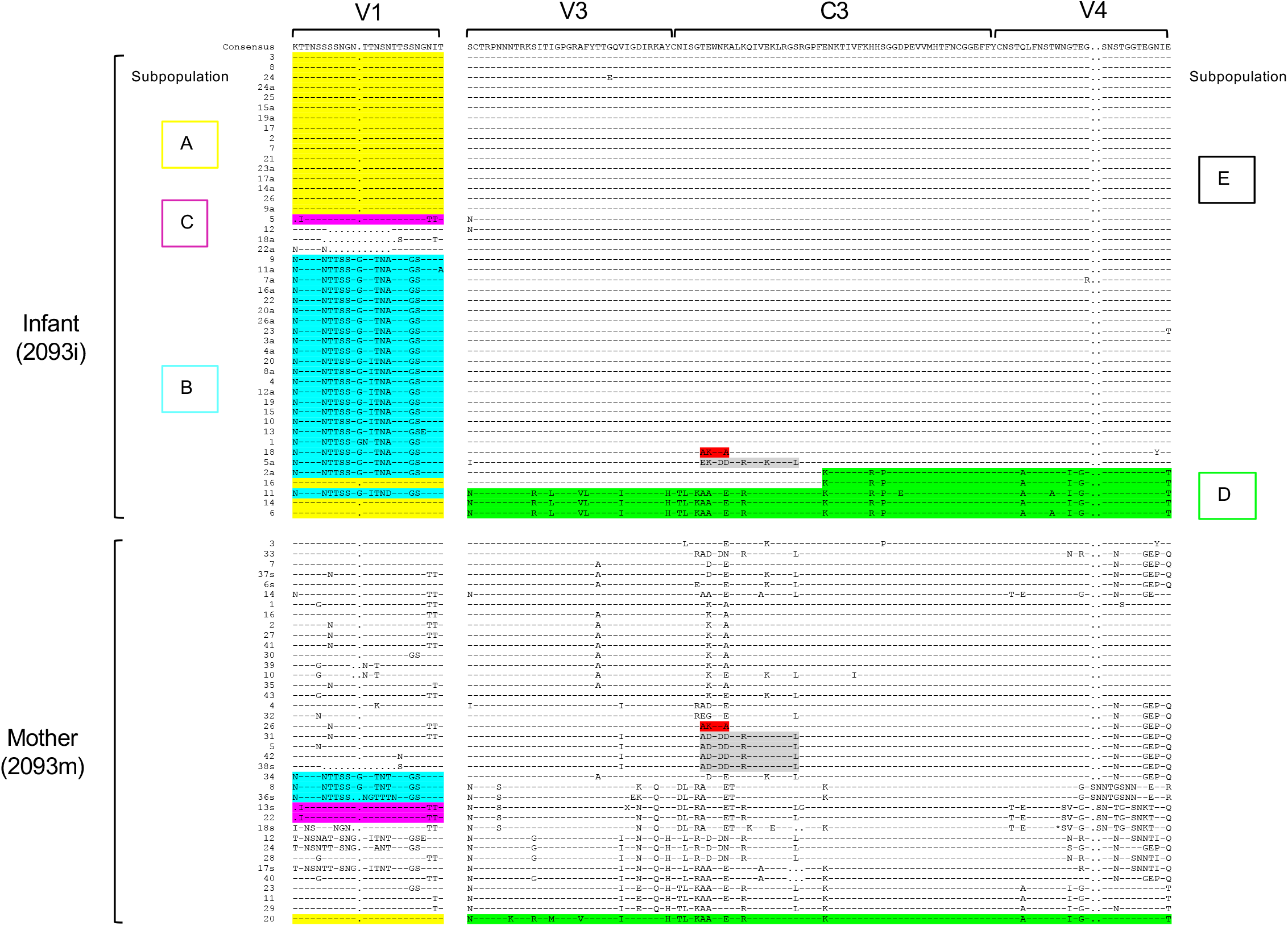
Identification of unique sequence motifs in the *env* gene from the mother-infant pair 2093. For each sequence, amino acid differences from the infant T/F virus are shown, while identical amino acids are represented as dashes. Distinct motifs shared between mother and infant (noted as A-E) are highlighted in different colors.

However, when looking at the two regions (V1 and V3C3V4) combined, the unique motif patterns observed in the infant sequences are all present in recombinant sequences as identified by RAPR analysis (**Fig. S2A**). Interestingly, no similar combinations of these motifs are found in the mother, indicating that these recombinants mostly likely generated in the infant after transmission. For example, recombinant sequences 1, 18, 5a and 2a all carry motif B in V1 and distinct motifs in the V3/C3/V4 region, and this pairing of the two motifs in the same sequence is not found in any of the maternal sequences (Fig. 3). All but two infant sequences (6 and 14) had pairings of unique motif patterns that were not present in the maternal viral sequences (**Fig. 3**). These results confirmed that the majority of the viruses in the long-term infected infants were recombinants as shown in **Fig. S2**. It is most likely that after the different viruses were transmitted from the mother to the fetus, these viruses recombined among them and only parts of transmitted maternal virus genomes were detectable later in the infant virus genomes. Similar results were also observed in 1580i (**Fig. S4**). Taken together, identifying origins of unique motif sequences of the infant *env* genes in their cognate mother virus populations shows that multiple variants are often transmitted into fetuses during long *in utero* infection time, and frequent recombination among them further increases the levels of genetic complexity of the viral population in fetuses.

### Host selection on viruses in fetuses

The high levels of genetic diversity in the viral populations in the four long-term infected infants (1039i, 2093i, 1580i and 2564i) prompted us to investigate whether specific loci within the *env* gene were under host selection pressure. We used the SNAP tool to determine the average pairwise cumulative codon behavior for synonymous (syn), nonsynonymous (non-syn) and insertions/deletion (indel) mutations across the entire *env* gene. For all four transmission pairs, nonsynonymous plots were higher than synonymous ones in both the infants and their cognate mothers (**Fig. 4**). The dS/dN (ω), the average ratio of the rate of nonsynonymous substitutions per nonsynonymous site, in infants 1039i and 1580i were 64% and 50% higher than their cognate mothers, respectively. However, overall there was no statistical difference between infants and mothers in the four pairs (p=0.38 by paired Wilcoxon test). Similar analyses of viral populations in the other eight infants who were infected for a short period of time showed similar accumulation of synonymous and non-synonymous mutations across the entire *env* gene in all but infant 1348i who was the oldest infants (40days) and had been infected for the longest time (61 days) among eight short-term infected infants (**Fig. S5**).

**Fig. 4.**
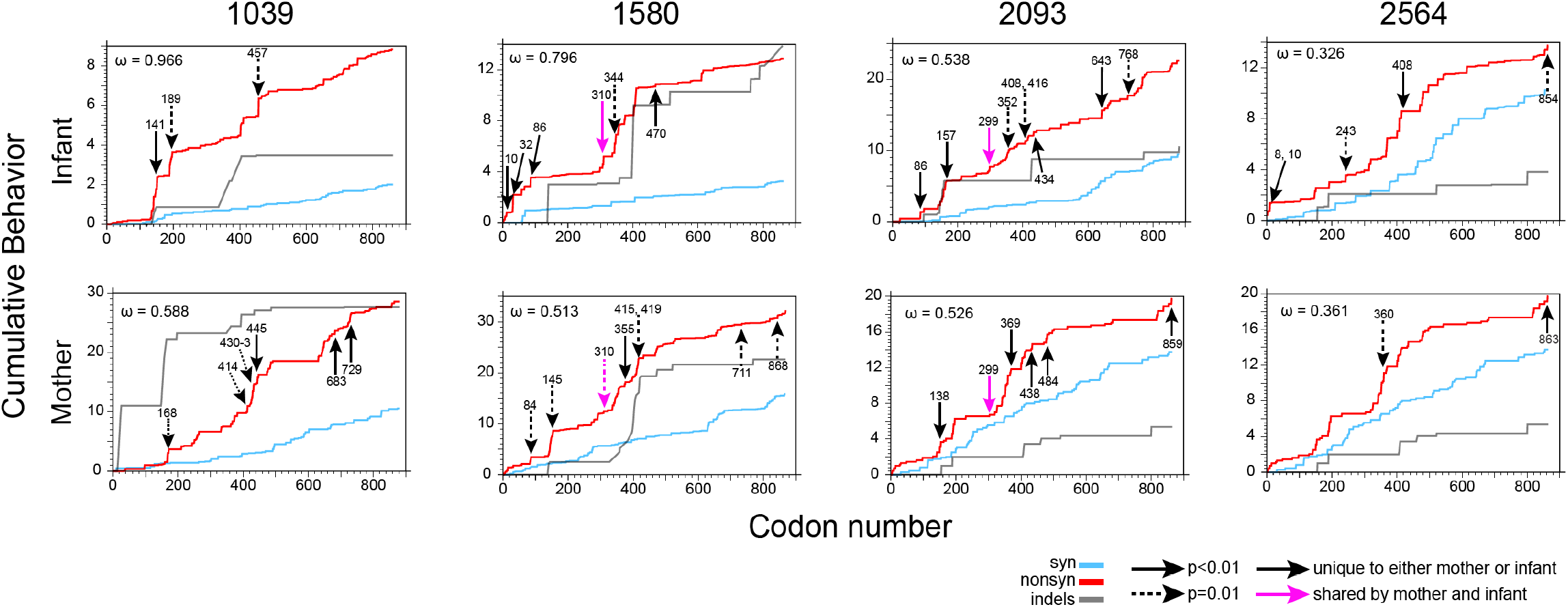
Identification of selection sties by analyzing the accumulation of mutations across the *env* gene in the infant viruses. Cumulative plots of each codon average behavior for all sequences pairs for the infant and maternal viruses for synonymous (green), non-synonymous (red) mutations and indels (blue). Sites found under statistically significant diversifying selection by the MEME analysis are marked by arrows, solid arrows indicate p<0.01 and dashed arrows p=0.01. Magenta arrows show sites under diversifying selection in both the mother and the infant. Values of ω denote average ratios of the rate of non-synonymous substitutions per non-synonymous site (dN/dS) for each sample.

In order to identify sites under positive diversifying selection, we analyzed the env sequences from the four long-term infected mother-infant pairs using the online tool MEME from the datamonkey server (Murrell et al., 2012). With a significance threshold of p=0.01 or lower, MEME identified a total of 23 sites significantly under diversifying selection among the four infants and 22 among the four mothers, with only two sites overlapping between mother and infant in two transmission pairs: at position 281 (based on the position in HXB2) in pair 2093 and position 304 in pair 1580 (**Fig. 4**). The former is one of the bindings sites for the broadly neutralizing antibody b12 (Zhou et al., 2007) and is associated to resistance to bNAbs VRC03, 4E10, and 2F5 (Bricault et al., 2019), while the latter site has been found to be associated with resistance to bNAbs PGT121 and PGT128 (Bricault et al., 2019). Positively selected sites were overwhelmingly concentrated in gp120 (20 out of 23 in the infants, and 17 out of 22 in the mothers), and, in particular, in the V3/C3/V4 region (14 out of 42 unique sites). Site 350 in particular, located in C3, a position significant in two mothers (1580m and 2093m, Fig. 4), is associated with resistance to bNAb CH102 and CD4 binding bNAbs (Bricault et al., 2019).

Examination of clustered mutations in the V1V2C2 region in 1039i (the first consecutive two vertical steeps of increases of non-synonymous mutations) showed that different regions were under disparate selection (**Fig. 5**). The C2 sequences were highly homogenous in 1039i and similar to half of those sequences in the mother. However, nearly half of the viral population in 1039m had variable C2 sequences. This suggests that the viral sequences in the region was highly selected in 1039i. In contrast, the V2 sequences were very homogenous in 1039m, except that one-third of the sequences had one A186T mutation, but about one-third of the sequences had various distinct mutations in 1039i. This suggests that this region was under positive selection in the infant, but not in the cognate mother. The V1 sequences were highly variable in both in 1039i and in 1039m. However, the sequences in the infant were different from those in the mother in this region (**Fig. 5**). Phylogenetic tree analysis confirmed that 101039i and 101039m sequences were distinct from each other (**Fig. S6**). Only one maternal virus sequence (101039m.20) clustered with two infant sequences (101039i.25 and 101039i.22). They differed from each other by one amino acid. These results indicate that V1 was under selection pressure in both 1039i and 1039m but the selection was different in the infant and the mother.

**Fig. 5.**
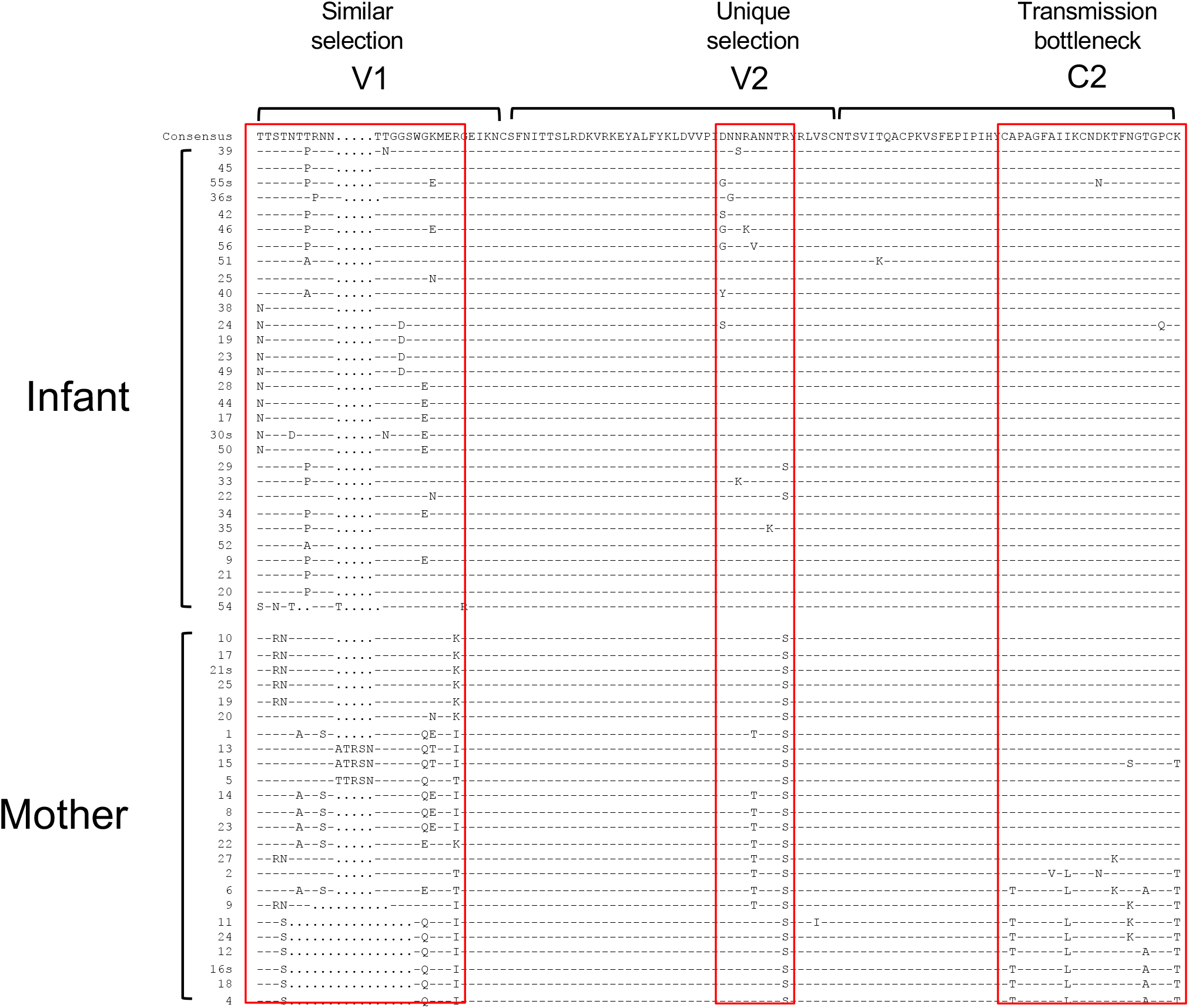
Selection signatures in *env* sequences from mother-infant pair 1039. Infant and mother sequences are compared to the infant consensus sequence. Distinct selection signatures at different sites in the V1V2C2 region in the infant and mother viruses are indicated by red boxes; similar selection in V1 for both infant and mother viruses, unique selection in V2 for the infant viruses, and transmission bottleneck (or purifying selection) in C2 for infant viruses.

Similar selection at the same region of the *env* gene in both infant and mother were also found for the transmission pairs 2093 and 2564. Divergent viral populations in both infants and mothers were detected in the cytoplasmic domain of gp41 for the pair 2093 (**Fig. S7A**) and in V5C5 for the transmission pair 2564 (**Fig. S7B**). New selection sites in the infant viral population were also detected in the V3 sequences in 2564i (**Fig. S7B**) and in the gp41 sequences in 1580i (**Fig. S7C**) while the viral populations were highly homogenous at the same region as the viruses in their cognate mothers. Taken together, detection of selection sites unique for infant viruses in all four long-term *in utero* infected infants indicates that viruses in fetuses are under host-specific selection pressure which are different from those in their cognate mothers.

Nine strong selection regions (**Figs. 5 and S7**) were identified among the four long-term *in utero* infected cases. Five of them were found in variable regions that are targeted by broadly neutralizing antibodies (Kwong and Mascola, 2018; Sok and Burton, 2018), indicating that selection pressure might be due to neutralizing antibodies (**Table S1**). The other four sites were found in the more conserved gp41 regions or the signal peptide where many T cell epitopes have been identified as reported in the HIV Immunology Database (http://www.hiv.lanl.gov/content/immunology), suggesting that these sites might have been targeted by T cell immune responses in fetuses (**Table S1**). These mutations were not random but rather concentrated in regions usually targeted by broadly neutralizing antibodies or T cell responses, suggesting that these sites were selected by host immune responses in the fetuses.

Since no viable cells were available from fetuses or infants from birth, we could not determine if these sites are targeted by the T cells in the infants. However, some residual plasma samples were available from both the infant and the mother form pair 1580 for us to determine autologous neutralization activity. We chemically synthesized two *env* genes that represent two sub-clusters in the infant viral population and one representative *env* gene in the maternal viral population and generated Env-pseudovirus for each *env* clone. Both infant pseudoviruses and the maternal pseudovirus were not or weakly neutralized by the infant or maternal plasmas (**Table S2**), indicating that both infant viruses are resistant to the plasma from both the infant or the mother.

### Higher neutralization activity in placenta plasma than systemic blood from the same HIV-1 infected mothers

To determine if viruses in the fetus may be exposed to differential neutralizing antibody pressure than those found in maternal plasma, we next sought to determine if neutralizing activities were different in plasma samples from systemic blood compared to placental blood of from an additional group of chronically-infected pregnant women. The plasma samples were collected from the placenta and peripheral blood of HIV-1 infected pregnant women. Interestingly, placental plasma neutralized tier-2 pseudoviruses TRO11 and BJOX2000 at higher titers than paired systemic blood plasma (**Fig. 6A**), although, placental plasma also showed higher non-specific antiviral properties against the pseudovirus generated with Env glycoprotein from murine leukemia virus (MuLV) than systemic blood plasma from the same mothers, indicating more non-specific neutralizing activity in placental plasma. However, neutralization titers of placental plasma were 2.97 to 11.98-fold higher than those of the systemic blood plasma from the same mothers. To determine if the neutralization activity was mediated by IgG, we depleted IgG from placental or systemic blood plasma samples and then determined their neutralization activity against TRO11. After IgG depletion, the neutralization activities were reduced to undetectable levels in over half of systemic and placental plasma samples (54%), or significantly reduced by 8.26 to 28.09 folds for the remaining samples (**Fig. 6B**), suggesting that IgG was responsible for the high levels of neutralization activity in placenta plasma. To determine whether the high titers of neutralization activities in placental plasma were due to high levels of HIV-1 Env-specific IgG, we compared binding Ab titers to two different HIV-1 gp120s (1086C and CON6) with paired systemic and placental plasma samples from the same mothers. The IgG titers to both 1086C and CON6 Env were similar between placental and systemic blood plasma from the same mothers (**Fig. 6C**). These results show that the IgG was responsible for the higher HIV-1 neutralization activity in the placental plasma compared to peripheral blood plasma from these mothers, indicating the potential for selective transfer of potently neutralizing IgG antibodies across the placenta (Martinez et al., 2019) and then contributes to selective pressure on fetal HIV variants.

**Fig. 6.**
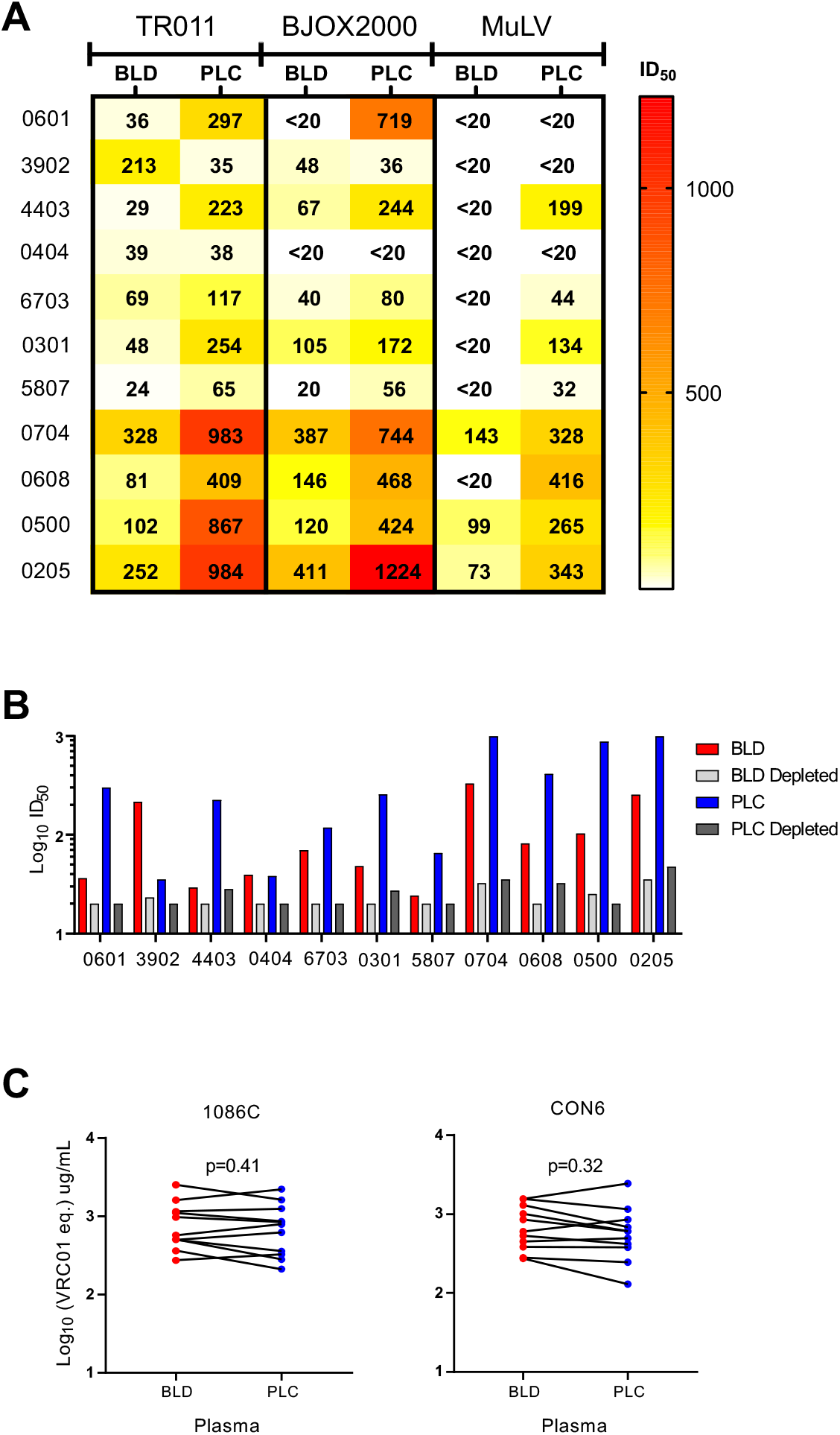
Higher neutralization activity of placental plasma than blood plasma from the same mothers. (A) Heatmap analysis of neutralizing titers of placental (PLC) plasma than systemic blood (BLD) plasma from the same mothers. The neutralization titers are color coded from dark to light where the darker color indicates higher neutralization titers. Patients in red are transmitting and those in black are nontransmitting. Asterisks indicate patients with placental plasma HIV-1 neutralization more than 3-fold higher than MuLV titers. BLD= Systemic blood plasma, PLC= Placental plasma. (B) Loss of neutralization activity after IgG depletion by protein G column in both placental plasma than systemic blood plasma. (C) Similar IgG concentrations in placental plasma than blood plasma from the same mothers. HIV-1 specific IgG concentrations were determined by measuring binding of blood and placental plasma to 1086C and CON6 gp120, respectively.

## Discussion

The placenta is a physical barrier which protects fetuses from infections by pathogens (Dauby and Marchant, 2020), but HIV-1 still can still pass through the placenta and infect fetuses in 5-10% of untreated HIV-1 pregnant women (Wolinsky et al., 1992). However, the time during pregnancy at which HIV-1 is more likely to cross the placenta and infect fetuses is not well understood. By analyzing the viral population using the SGA method, we found that T/F viruses could be well defined in eight out of 12 *in utero* infected infants. The median of days of infection was 18 (3-37) days before birth. The majority (67%) of *in utero* infections happened within the last 40 days before delivery during pregnancy in this cohort. The placental barrier changes dynamically during pregnancy. In the first trimester, it consists of the syncytiotrophoblast, the cytotrophoblast, the villus mesenchyma, and the fetal capillary walls. During the last trimester, the thickness of the barrier significantly decreases as the cytotrophoblast disappears and fetal vessels get closer to the villus surface. The exchange zones of the barrier consist of only membranes of syncytiotropholast and adjacent capillaries walls (Chatuphonprasert et al., 2018; Codaccioni et al., 2019; Schatten and Constantinescu, 2007). Although it is unclear how HIV-1 viruses cross the placenta, the risk of transmission is increased during the third trimester when the placental barrier becomes thinner. This is consistent with our finding that most of *in utero* MTCT occur during the late stage of pregnancy.

Conventional T lymphocytes can be detected in fetal spleen and lymph nodes only by 14 weeks of gestation (Dauby and Marchant, 2020). Thus, HIV-1 infection may be less likely to occur in fetuses during early pregnancy when T cells have not developed yet. All distinct motifs in the infant viral populations can be identified in the viral sequences in their cognate mothers. Thus, they are most likely the result of recombination among transmitted maternal viruses carrying these motifs and selection in the infants. The current molecular clock analyses which assume no recombination in the analyzed sequences puts the infection time way before the time when T cells are detected in fetuses (14 weeks of gestation) or even before the pregnancy starts. Our study demonstrated that recombination among viruses in fetuses is extensive and that further confirms such sequences cannot be used to reliably estimate the infection time (Song et al., 2018).

Three out of eight (37%) *in utero* infections for which T/F lineages could be defined were infected by more than two T/F viruses. The rate in this small sampling of *in utero* infections is nearly two times higher than the rate (20%) among heterosexually infected people (Abrahams et al., 2009; Keele et al., 2008) and similar to the rate (36%) among men who having sex with men (MSM) in a subtype B infection cohort (Li et al., 2010). Since infection with more than one T/F viruses is associated with higher risk for HIV-1 transmission (Bar et al., 2010; Li et al., 2010; Santra et al., 2015), our data suggest that the MTCT risk of HIV-1 infection during late pregnancy in the absence of maternal antiretroviral therapy may be similar to that in the MSM population.

The genetic diversity levels of the viral populations in infants with long-term *in utero* infection was very high, only about three folds lower than those in their cognate mothers, while the difference was as high as 13 folds lower when the genetic diversity in the other eight infants infected only for a short time were compared to their mothers. The diversity of the viruses in infant 2564i was only slightly lower than that of the cognate maternal viruses. Although the genetic diversity levels were high in the infant viral populations, the infant viruses formed independent lineages that are separate from their cognate maternal viruses, strongly suggesting that the infant viruses have different evolutionary pathways from the viruses in the cognate mothers during *in utero* infection. The only exception is that a few viruses in 2093i formed two sub-clusters with the maternal viral sequences while the majority of the infant viral sequences formed an independent cluster. Further analysis showed that those infant sequences clustering with the mother viral sequences are recombinants.

Sequence analysis showed that many *env* gene regions were highly homogenous in four long-term *in utero* infected infants while the sequences were highly divergent at the same region in their cognate mothers, suggesting a transmission bottleneck. However, infant viral sequences were divergent at some regions of the *env* gene where the cognate maternal sequences were homogenous or similarly divergent but with different sequences. These results indicate that the viruses are under different selection pressures in the fetuses compared to the maternal viruses.

Although fetuses are generally thought to reside in a mostly sterile condition, they can be exposed to a wide range of immuno-stimulatory molecules such as ingested amniotic-fluid substances (Mold et al., 2008; Underwood et al., 2005), semi-allogenic antigens from maternal cells (Aagaard et al., 2014; Claas et al., 1988), microbes (Campbell et al., 2015), and food antigens (Berry et al., 1992). Innate immune responses and antigen-specific T cell responses to pathogens are detectable in fetuses, suggesting that the exposure to maternal infection *in utero* may prime the developing immune system (Wilcox and Jones, 2018). Aberrant fetal immune responses to maternal antigens is found to play an important role in the pathogenesis of preterm labor (Frascoli et al., 2018). A recent study showed that as early as the second trimester fetal antigen presenting cell (APC) subset such as conventional dendritic cell (DC) can respond to toll-like receptors (TLR) (McGovern et al., 2017). However, when compared to infant immune responses, fetus immune responses mediated by macrophages are more geared towards immune tolerance. Precursors of mature B lymphocytes (pre-B cells) are detected in the fetal bone marrow by 13 weeks of gestation (Dauby and Marchant, 2020). Although the diversity of the immunoglobulin repertoire is limited during the initial stages of B-cell development but appears to be similar to that of adults by the third trimester of gestation (Rechavi et al., 2015). However, the role of antibody responses in preventing *in utero* HIV-1 infections is still unclear (Dauby and Marchant, 2020; Douglas et al., 2017).

HIV-1-specific CD8+ T-cell activity is detectable from birth in up to 70% of *in utero* infected infants (Thobakgale et al., 2007), suggesting that T cell responses may exert selection pressure on HIV-1. Fetal exposure to HIV-1 primes the immune system to enhance immune activation and alters T cell homing in fetuses (Bunders et al., 2014). Although these studies suggest immune responses in fetuses, no data shows that adaptive immune responses play a role in shaping pathogen evolution. In this study, we show that more than half of the selected sites in the *env* gene in fetuses were found in variable regions targeted by broadly neutralizing antibodies (Kwong and Mascola, 2018; Sok and Burton, 2018), indicating selection pressure possibly triggered by neutralizing antibodies. The other half of the sites were found in the conserved gp41 regions or in the signal peptide, both of which are frequently targeted by T cell responses (http://www.hiv.lanl.gov/content/immunology). Because there were no cells and sufficient amount of plasma samples available from these studied subjects, we could not determine HLA alleles in these transmission pairs or whether those sites in the *env* gene are likely selected by T cell responses. However, the detection of different amino acids in the infant viruses at the previously well-defined T cell epitopes compared to the cognate maternal viruses indicates that T cell responses may also play a role in selecting these mutations. Taken together, these data suggest that host selection pressure has played an important role in selecting escape mutations and shaping the viral evolutionary pathway in fetuses.

The immune responses in fetuses are difficult to study because it is nearly impossible to obtain adequate blood samples from fetuses during pregnancy. In addition, maternal antibodies and half of HLA alleles are transferred from the mother to the fetus, making it a challenge to determine whether the immune responses are from the infected fetuses or their cognate mothers. However, the detection of stronger IgG mediated neutralization activity from placental plasma compared to the systemic plasma in the same mothers at birth indicates that placental selection of antibodies for transfer to the infants may play a role in blocking MTCT transmission and driving these mutations in variable loops in the *env* gene. It is also plausible that paternal HLA alleles may contribute to the selection of mutations at those well-defined T cell epitope sites, which are not likely to be associated with neutralizing antibodies. The presence of distinct mutation clusters in the *env* gene of infant viruses compared to the same sites in cognate mother viruses clearly shows that the selection at these sites is due to unique immune response in fetuses.

The first samples in all four long-term *in utero* infection were collected between one and two months after birth. To investigate if the high levels of genetic variation and selection mutations were generated during the infection period after birth, we determined how many mutations can accumulate within 1-2 months after birth in four infants for whom subsequent samples were available for analysis. Since the immune responses are expected to be more mature and potent in infants after birth, we would expect to see more selected mutations in the viral genome after additional infection time. However, no obvious additional mutations were detected in all four infant viruses even three more months after the first analyzed sample. Second, when we compared the viral populations from two short-term and two long-term infection cases with similar time after birth, we found that long-term infection infants had more genetically diverse viral populations, while the short-term infection cases had more homogenous viral populations, with only a few single base selection sites, which is typically observed as in our previous analysis of acute/early infection during the first 2 months of infection in adults (Gao et al., 2014; Liu et al., 2013; Ritchie et al., 2014). Thus, our results demonstrate that diversity in these long-term *in utero* infections was driven by recombination and new mutations mainly accumulated during pregnancy, not after birth.

In summary, we performed detailed genetic analyses of HIV-1 *env* gene sequences from perinatal transmission pairs and found that most *in utero* transmissions occur just before delivery in the third trimester. Moreover, host selection can be detected during long-term *in utero* infection of the fetuses. Knowing the timing of *in utero* infection during pregnancy will be important for the development of more effective approaches to prevent perinatal MTCT. The identification of strong selection sites potentially mediated by neutralizing Abs and T cell immune responses indicates that the immune system in the fetus is capable of exerting selection pressure on viral evolution during *in utero* infection. A similar approach can also be applied to other viral infections. Studying the selection and evolution of HIV-1 and other viruses in fetuses can be an ideal alternative approach to study the development of adaptive immunity in fetuses.

## Acknowledgments

We would like to acknowledge the support of Pediatric HIV/AIDS Cohort Study (PHACS) team for their management of the Women and Infant Transmission study (WITS) cohort samples (U01 HD052102-04 and U01 HD052104-01), and the CHAVI009 team, and all study participants. David R. Martinez was supported by an American Society for Microbiology Robert D. Watkins Graduate Research Fellowship, a Burroughs Wellcome Graduate Diversity Fellowship, and an NIH National Institute of Allergy and Infectious Diseases (NIAID) Ruth L. Kirschstein National Research Service Award F31 F31AI127303. This work was supported by a grant (1R01AI22909) from National Institutes of Health/ National Institute of Allergy and Infectious Diseases (NIH/NIAID).

## Author contributions

Conceptualization, F.G. and S.R.P; Data collection, M.H.M, M.M., A.K., J.J.T., D.R.M. and X.L.; Data analysis, M.H.M, M.M., A.K., E.E.G., J.J.T., L.F., S.R.P and F.G.; Writing – Original Draft, M.H.M. and F.G.; Writing – Review & Editing, M.H.M, M.M., A.K., J.J.T., D.R.M., E.E.G., L.F., S.R.P and F.G.; Funding Acquisition, S.R.P and F.G.

## Declaration of interests

The authors declare no competing interests.

## Notes

### Competing Interest Statement

The authors have declared no competing interest.

