## Supplemental files for "Different evolutionary pathways of HIV-1 between fetus and mother perinatal transmission pairs indicate unique immune selection pressure in fetuses"

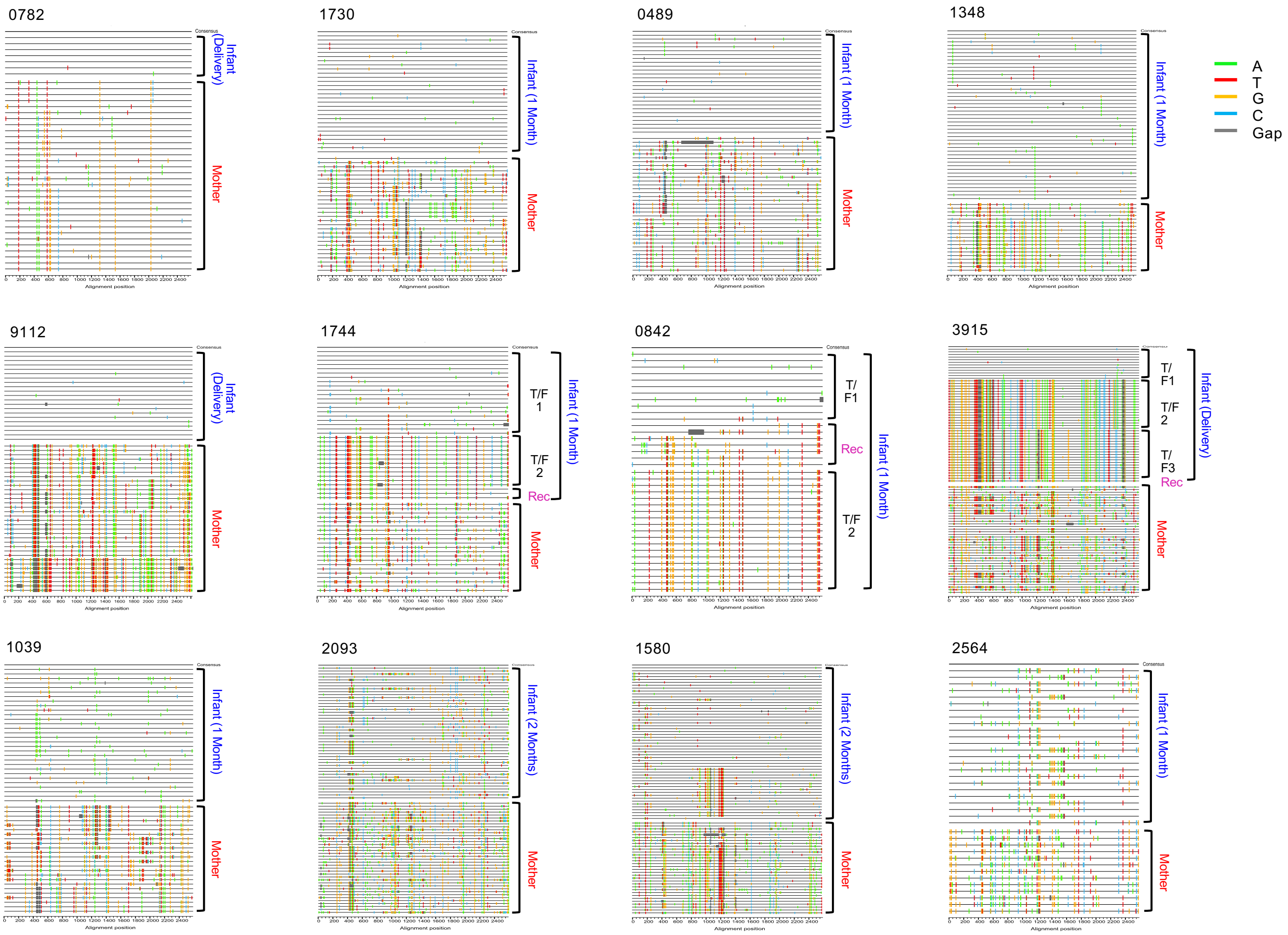


**Figure S1. Highlighter plots of the env sequences from each mother-infant transmission pair.** Highlighter plots show the location of all nucleotide substitutions in each SGA-derived *env* sequence compared to the infant consensus sequence shown at the top. When more than one T/F viruses are present, the T/F from the major lineage (T/F1) is used as reference. Each line represents one *env* sequence. The positions of these substitutions are indicated on the bottom axis. Nucleotide substitutions and gaps are color coded. T/F sequences are inferred for the top eight infants, while they cannot be inferred for the bottom four infants due to the long-term infection during pregnancy. Recombinant sequences (Rec) among infant viruses are indicated in magenta.

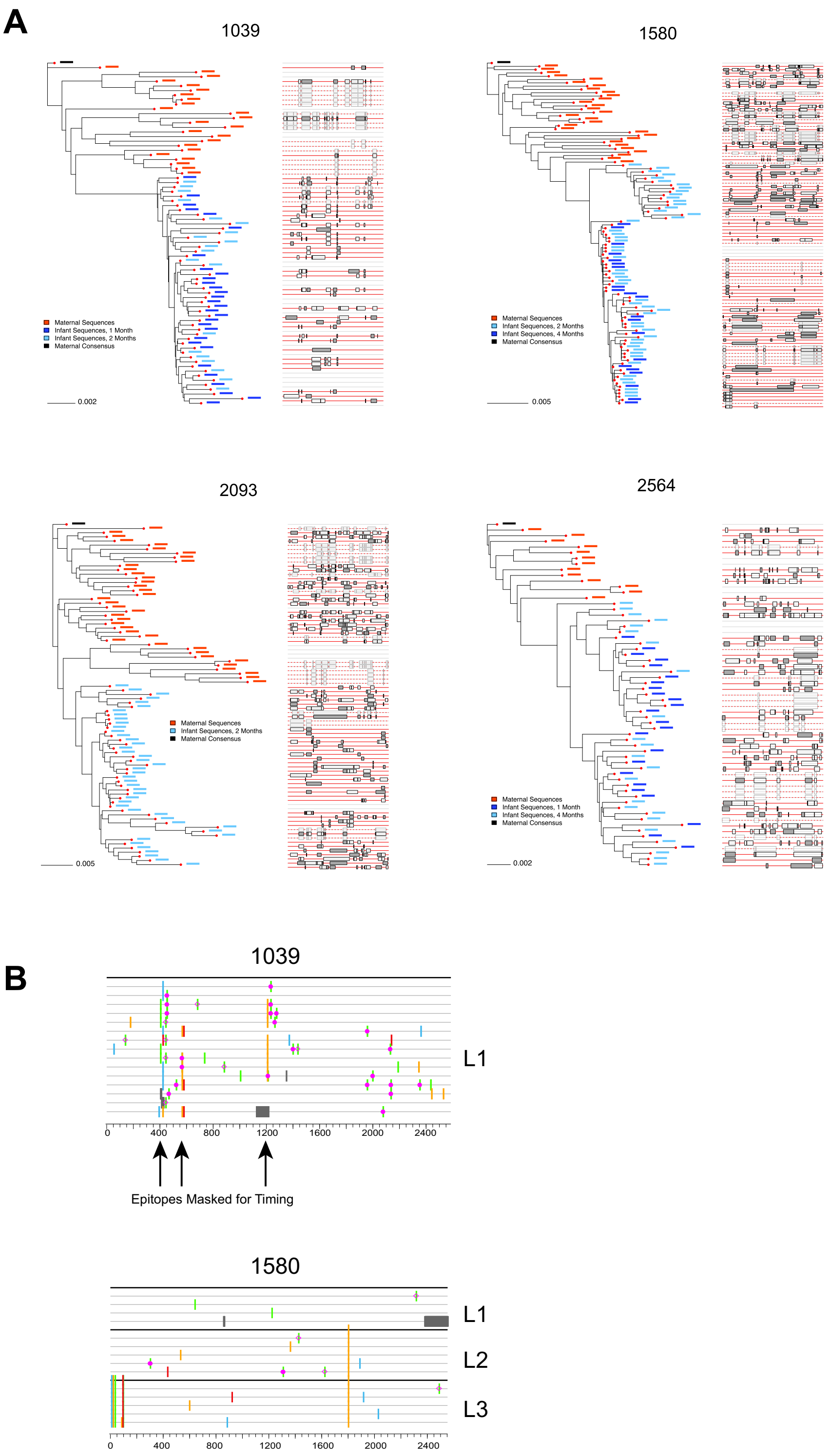


**Figure S2. Extensive recombinant among viral populations in infants and mothers after long-term infection.** (A) Phylogenetic tree of the env sequences from each transmission pair and recombination breakpoints as determined by the RAPR tool for mother-infant transmission pairs 2093, 1039, 1580 and 2564. Each mother-infant sequence alignment was analyzed through the recombination detection tool RAPR using the maternal sequence consensus as reference. In the phylogenetic tree, red bars indicate infant sequences and blue bars indicates maternal sequences. In the recombination breakpoint plot on the right, each line represents one sequence, and red colored lines represent the recombinants. Solid lines represent independent recombinant, while dashed line represent sequences that belong to a recombinant lineage. The range intervals where breakpoints are most likely to occur are indicated by black boxes, and boxes shaded in gray denote statistically significant breakpoints. (B) Highlighter plots of the lineages extrapolated from the two long-term infected infants after removing recombinants. Gray lines represent sequences and colored tic marks represent mutations from the consensus (top sequence in black). All recombinants detected by RAPR were removed with the exception of recombinants that gave rise to lineages with 4 or more sequences. Each lineage consensus is shown in black.

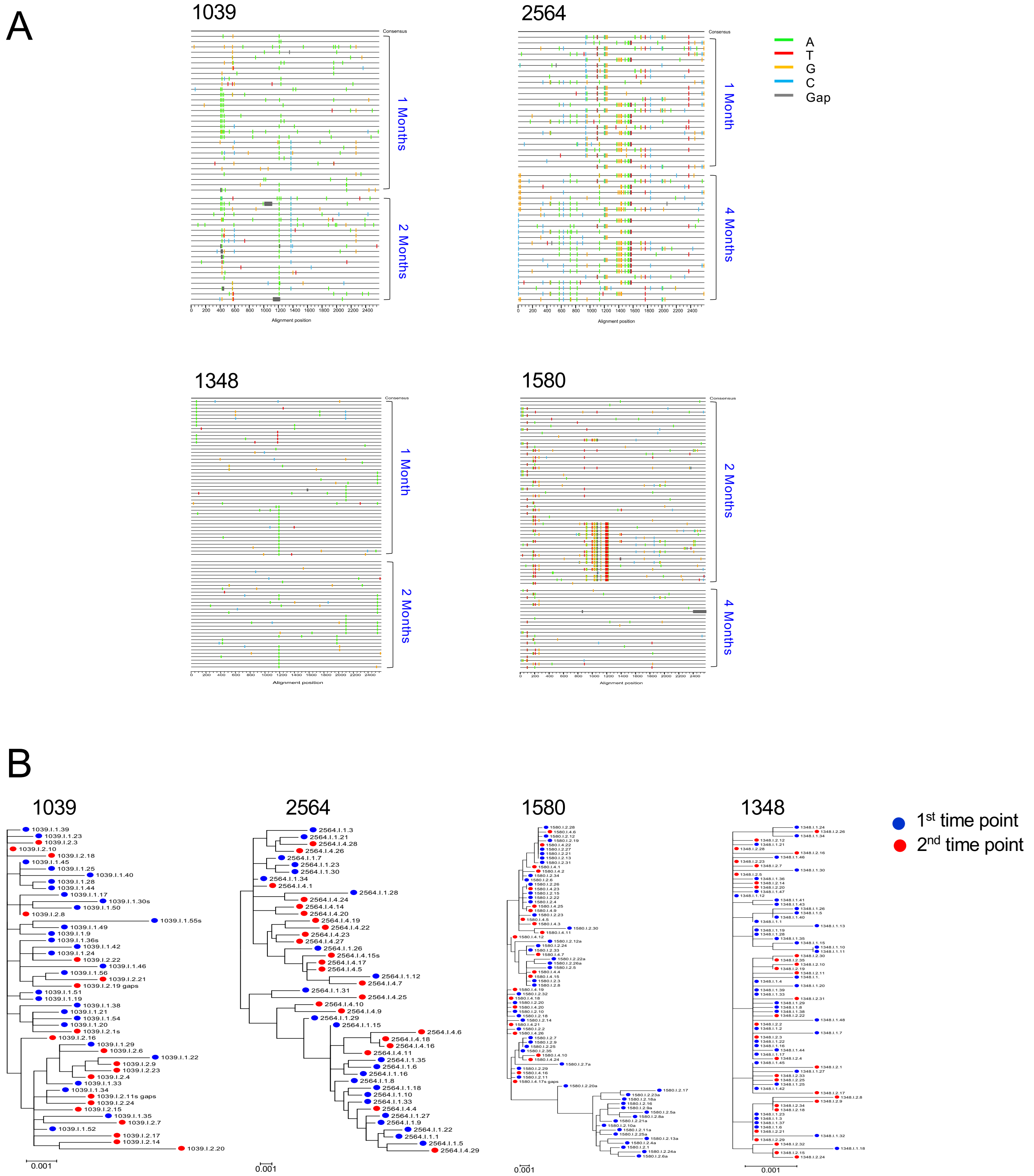


**Figure S3. Highlighter plots and phylogenetic analysis of infant *env* sequences from two time points.** (A) Highlighter plot denotes the location of nucleotide substitutions in each SGA-derived *env* sequence compared to the consensus or T/F1 (when more than one T/F viruses are present) sequences of viruses from infants. Each line represents one *env* sequence. The positions of these substitutions are indicated on the bottom. Nucleotide substitutions and gaps are color coded. No obvious differences are noticed in viral populations from two different time points. The only exception is that a subpopulation at month 2 is not detectable at month 4 in 1580. (B) Phylogenetic trees of the infant *env* sequences from two time points. Phylogenetic trees were constructed by the neighbor-joining method with the Kimura 2-parameter model. Sequences from two two different time points are indicated in blue and red dots, respectively. The viral sequences from two sequential time points from the same infant are phylogenetically indistinguishable.

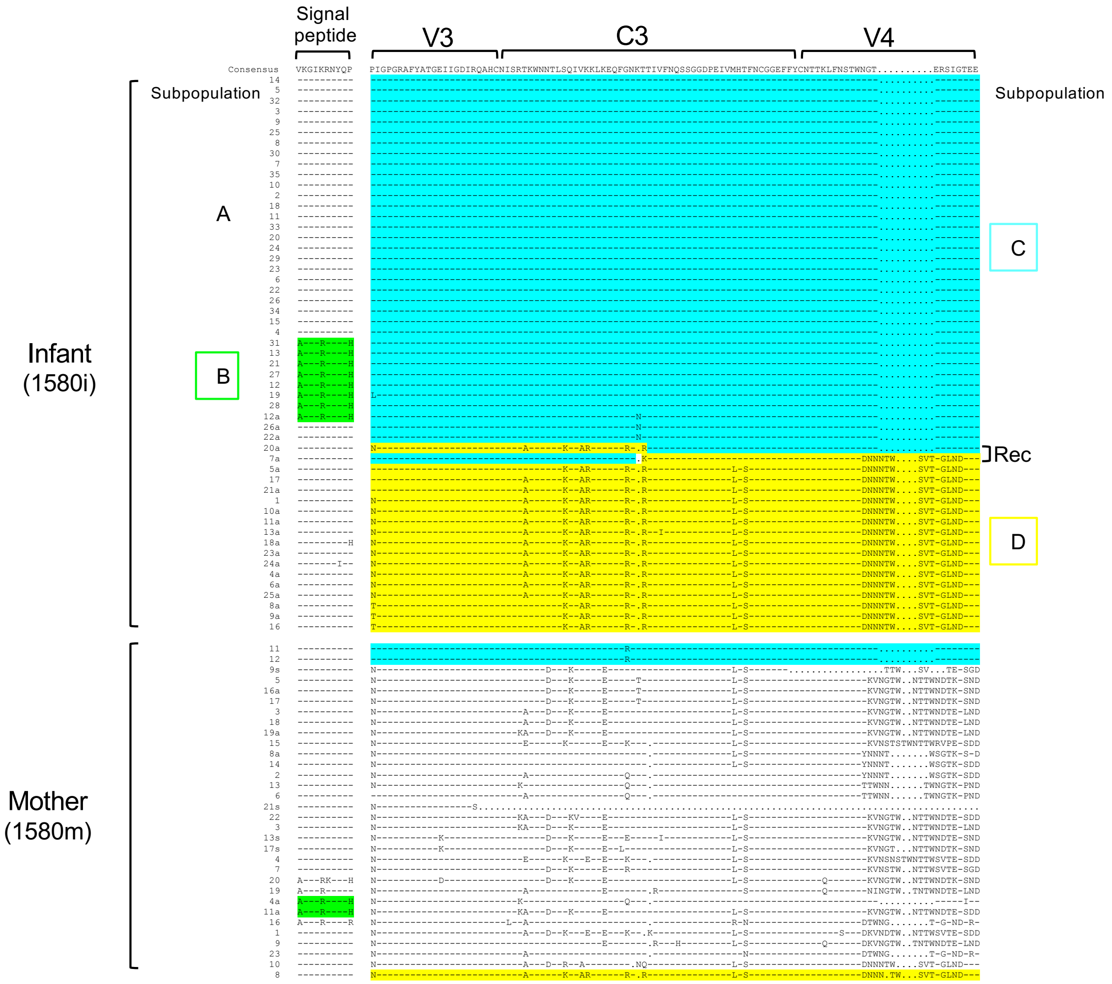


**Figure S4. Identification of unique sequence motifs of infant viral sequences in the cognate mother viral sequences.** The infant and mother sequences are compared to the consensus sequences of the viruses in infant 1580i. Distinct motif sequences in the fetal viral population are indicated by different colors, while the identical or similar motif sequences found in the maternal virus sequences are indicated with the same colors. Two recombinant sequences (Rec) in the infant are indicated.

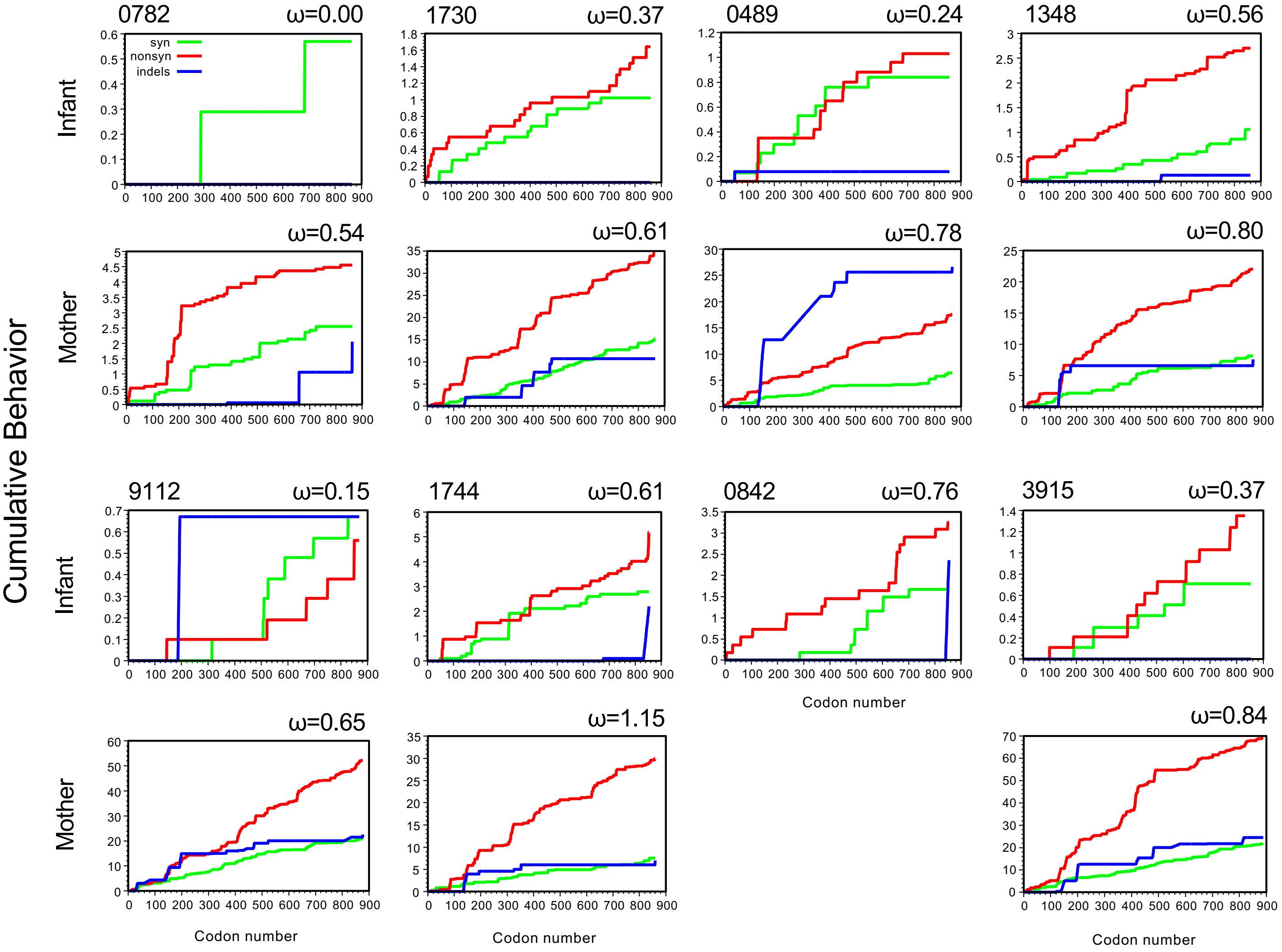


**Figure S5. Accumulation of mutations across the entire *env* gene in the infant and mother viruses.** Cumulative plots of each codon average behavior for all sequences pairwise comparison of infant or mother viruses for synonymous (green), non-synonymous (red) mutations and indels (blue). Values of ω denote average ratios of the rate of nonsynonymous substitutions per nonsynonymous site (dN/dS) for each virus.

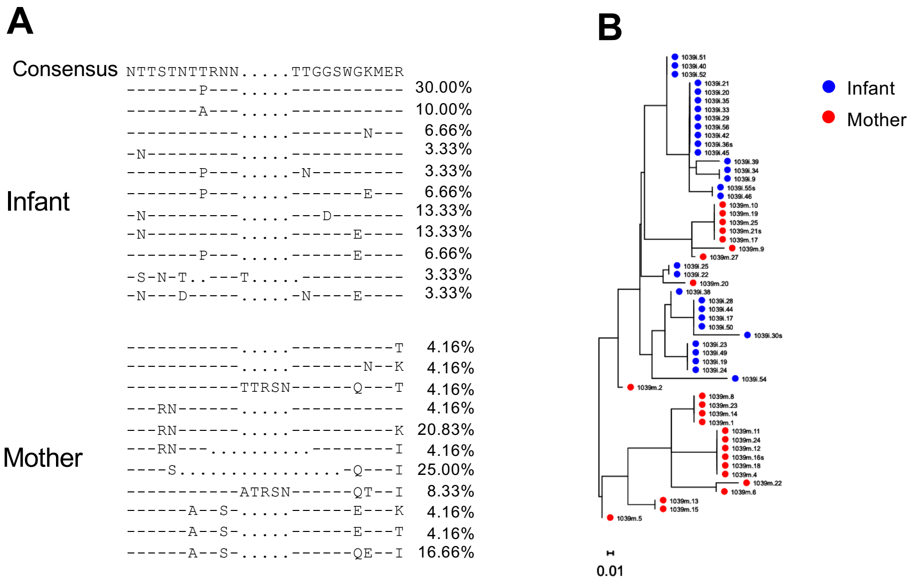


**Figure S6. Distinct amino acid signature patterns in the infant and mother viruses in transmission pair 1039.** The V1 sequences are highly divergent in the infant and mother viruses (A). Only a few sequence patterns account for more than 20% of the viral population in infant or mother. Phylogenetic tree shows that no infant viral sequences are identical to the mother viral sequences (B). Only two infant viral sequences are similar to one mother viral sequence.

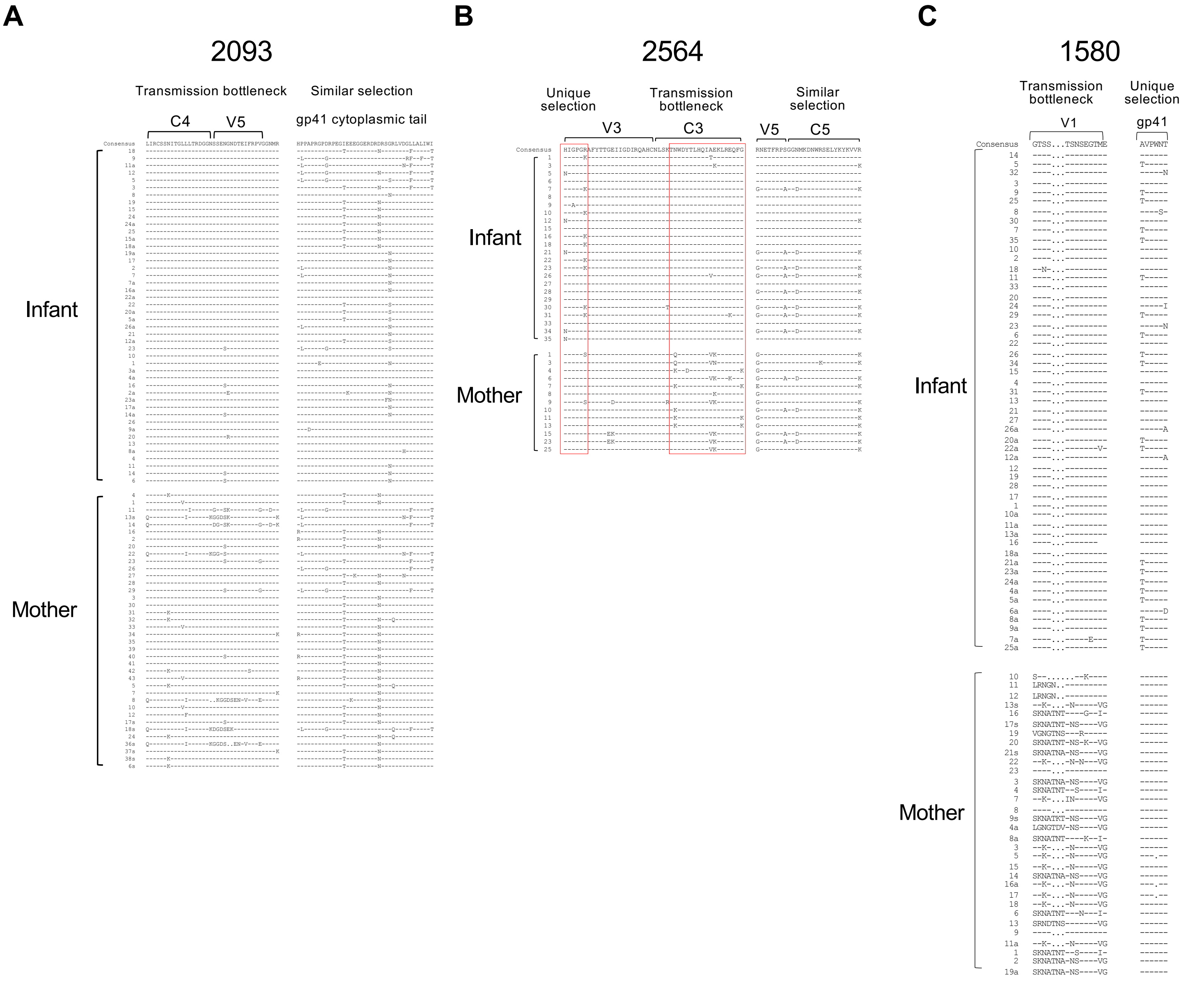


**Figure S7. Selection signatures in the *env* sequences in the infants**. (A) The infant and mother sequences are compared to the consensus sequences of the viruses in infant 1039i. Distinct selection signatures at different sites in the C4V5 region and the cytoplasmic tail of gp41 in the infant and mother viruses are shown; similar selection in the gp41 cytoplasmic tail for both infant and mother viruses, and transmission bottleneck in C4V5 for infant viruses. (B) The infant and mother sequences are compared to the consensus sequences of the viruses in infant 2564i. Distinct selection signatures at different sites in the V3C3 region in the infant and mother viruses are indicated by red boxes; unique selection in V2 for the infant viruses and transmission bottleneck in C3 for infant viruses. Similar selection is detected in V5C5 for both infant and mother viruses. (C) The infant and mother sequences are compared to the consensus sequences of the viruses in infant 1580i. Distinct selection signatures at different sites in V1 and gp41 in the infant and mother viruses are shown; unique selection in gp41 for the infant viruses and transmission bottleneck in V1 for infant viruses.

**Table S1. Potential selection sites by CTL and neutralizing antibodies in infant viruses.**

| Virus | Selection site | Selection region | Variation in known CTL epitopes |
|---|---|---|---|
| 1039i | #1 | V1-V2 | No |
| | #2 | V4-V5 | No |
| 2093i | #1 | V1 | No |
| | #2 | V3-V4 | Yes |
| | #3 | gp41 (around TMD) | Yes |
| 1580i | #1 | Signal peptide, C1 | Yes |
| | #2 | gp41 (before TMD) | Yes |
| 2564i | #1 | C3-V4 | Yes |
| | #2 | gp41 (N-terminus) | Yes |

ND: not detected, TMD: transmembrane domain

**Table S2. Neutralization susceptibility of infant and mother pseudoviruses to autologous plasma from pair 1580.**

| Virus | Infant plasma $ID_{50}$ (dilution) | Maternal plasma $ID_{50}$ (dilution) | CH103 $IC_{50}$ (μg/mL) |
|---|---|---|---|
| Infant 1 | <40 | <40 | 0.69 |
| Infant 2 | 42 | <40 | 1.13 |
| Mother | <40 | <40 | 1.13 |
| SF162 (tier 1) | 55 | 45 | 0.17 |
| TR011 (tier 2) | <40 | 52 | 5.76 |
